# Three-dimensional culture of chicken primordial germ cells in chemically defined media containing the functional polymer FP003

**DOI:** 10.1101/358952

**Authors:** Yi-Chen Chen, Wei-Che Chang, Shau-Ping Lin, Masataka Minami, Christian Jean, Hisato Hayashi, Sylvie Rival-Gervier, Tatsuro Kanaki, Shinn-Chih Wu, Bertrand Pain

## Abstract

Scalable production of avian suspension cell exhibits a valuable potential on therapeutic application by producing recombinant protein and as the substrate for virus growth. This study sought to establish a system with chemically defined components for three-dimensional (3D) culture of chicken primordial germ cells (cPGCs), a pluripotent avian cell type. cPGCs were cultured in medium supplemented with the functional polymer FP003. Viscoelasticity was low in this medium, and cPGCs did not sediment,and consequently their expansion was improved. The total number of cPGCs increased by 17-fold after 1week of culture in 3D-FAot medium, an aseric chemically defined medium containing FP003, indicating that this medium enhances the expansion of cPGCs. Moreover, cPGC cell lines stably expressed the germline-specific reporter VASA:tdTOMATO, as well as other markers of cPGCs, for more than 1 month upon culture in 3D-FAot medium, indicating that the characteristics of these cells are maintained. cPGCs harboring both PGK:EGFP and VASA:tdTOMATO robustly expressed both fluorescent proteins upon culture in 3D-FAot, suggesting that this approach is perspective for recombinant protein production. In summary,this novel 3D culture system can be used to efficiently expand cPGCs in suspension without mechanical stirring or loss of cellular properties. This system provides a platform for large-scale culture ofcPGCs in industry.

## Introduction

A three-dimensional (3D) sphere cell culture system that does not require mechanical stirring or agitation was established using the properties of a polysaccharide polymer [1]. Moreover, 3D culture systems enable mammalian and human embryonic stem cells, induced pluripotent stem cells, and hepatocytes derived from these cells to float in the culture medium [2–4]. In traditional medium, cells eventually settle on the bottom of the culture dish due to the effect of gravity and may subsequently lose critical properties. Preventing cells attaching to the surface of the dish would help to overcome such problems and enable efficient utilization of culture space and resources. In addition, cells can be cultured on a large scale using 3D systems. Suspension cells could be potentially cultured in large-volume bioreactors using 3D culture medium to produce a large number of cells for industrial manufacture of recombinant proteins.

Recombinant proteins have many therapeutic purposes, and consequently severalsystems have been established for their industrial production. *Escherichia coli* has been used to produce recombinant proteins because it can be easily cultured and is amenable to genetic modification. However, the production of recombinant proteins using this system is hampered by a lack of post-translational modifications (PTMs) and the risk of endotoxin contamination [5]. Recombinant proteins are also frequently produced in yeasts, such as *Saccharomyces cerevisiae* and *Pichia pastoris*. Although yeast cells can be easily and inexpensively cultured, this approach is restricted by the limited number of yeast vectors and promoters as well as the lack of PTMs observed in human cells [5, 6]. Production of recombinant proteins in animal cells is a promising alternative with various clinical applications. Animal cells need to be cultured for longer than microbial cells, and their culture conditions are more complicated. However, proteins produced in animal cells are similar to human proteins in terms of their PTMs (including glycosylation) and folding. Therefore, therapeutic proteins, especially those with complex structures, monoclonal antibodies (mAbs) as an example, are predominantly produced in Chinese hamster ovary cells [7–9]. Carbohydrate moieties of antibodies play a crucial role in the efficacy of antibody-based therapies [10]. Removal of fucose from IgG1 oligosaccharide improves binding to Fcg receptor IIIa on effector cells [11]. However, it is difficult to produce large amounts of non-fucosylated therapeutic mAbs from mammalian cells. Avian species can produce glycoproteins with a low level of fucosylation, which enhances antibody-dependent cellular cytotoxicity [12, 13]. In addition, species-specific glycosylation of recombinant proteins in host cells may pose a risk to human health due to the potential of these proteins to induce immunogenicity. N-glycolylneuraminic acid is attached to the terminal N-glycan of most proteins produced in mammals. This moiety is not found in humans and has a high potential to trigger allergic reactions [14]. Fortunately, N-glycolylneuraminic acid is not present in chickens. In addition, the humanized glycan N-acetylneuraminic acid is added to the terminal residue of N-glycans in chickens [14]. Moreover, the N-glycan profile of chicken IgY is reportedly suitable for the production of therapeutic mAbs [15]. Chicken cells have the potential to produce high-quality mAbs, and consequently may also be suitable for generating functional peptides.

Transgenic hen as bioreactor to produce therapeutic protein in laid egg has been produced and some of the lines show high to low productivity of the proteins of interest [16, 17]. However, it is tedious to generate each transgenic chicken line and to select strains with the better productive efficiency. In addition, the safety risk of those in vivo transgenic chicken will always be a troublesome problem and could be a limitation for a large pharmaceutical interest and manufacturing those products even if the recombinant protein purification from egg white is usually well achievable. Therefore, cell-base bioreactor becomes an alternative for the purpose of pharmaceutical protein production. Though oviduct epithelial cells show the application potential [18, 19], the absence of established lines and the limited number of passages of those primary adherent cell types are major blockages for a large scale industrial production. Thereafter, avian pluripotent cell displays the ideal model for this purpose. For example, EB66 cell line derived from duck embryonic stem cells exhibits efficient productivity in therapeutic monoclonal antibodies [12]. For the industrial interesting application, EB66 and other avian cell lines, e.g. AGE1.CR and QOR2/2E1, can also be adapted to cell suspension culture, which allows a scalable production in a large-scale bioreactor. Therefore, these cell lines in industry also progressively replace the primary cells to become cell substrates for virus replication for the vaccine production [20, 21].

Among the avian stem cells, chicken primordial germ cells (cPGCs) are germline stem cells taken from the embryonic blood in the dorsal aorta of HH15–16 embryos and exhibit in *in vitro* culture long term self-renewal potential [22–24]. Moreover, cPGCs are cultured in suspension and in an anchorage-independent manner, but no report was yet available as a large-scale production. Therefore, the present study aimed first to establish a 3D suspension culture system for cPGCs using chemically defined media containing a functional polymer FP003, second to characterize the lines grown in those conditions by specific markers and developmental potential and finally to demonstrate that the 3D suspension culture system allows the production of ectopic fluorescent protein expression. As a perspective, this study demonstrates that the cPGCs can be cultured in large bioreactors and could be useful for therapeutic protein productions

## Materials and methods

### Incubation of chicken eggs

To isolate cPGCs, specific pathogen-free chicken (*Gallus gallus*) eggs were purchased from the Animal Drugs Inspection Branch, Animal Health Research Institute, Council of Agriculture, Executive Yuan, Taiwan. All chicken embryos were cultured in a humidified incubator at 37.5 °C and automatically turned. All animal experiments were conducted with the ethical approval of the Ilan Branch of the Taiwan Livestock Research Institute (No. 105-11).

### Preparation of culture media

The three types of culture media used in this study were prepared as described by Whyte et al. with minor modifications [24]. FAcs medium was diluted DMEM (1:3 ratio of sterile water:calcium-free DMEM) containing 1× B-27 supplement, 2 mM GlutaMax, 1× non-essential amino acids, 0.1 mM β-mercaptoethanol, 1 mM sodium pyruvate, 0.2 % chicken serum (all purchased from Gibco®, USA), 1× nucleosides (EMD Millipore, USA), 2 mg/mL ovalbumin (Sigma-Aldrich, Germany), 0.1 mg/mL sodium heparin (Sigma-Aldrich), 25 ng/mL human Activin A, and 4 ng/mL human fibroblast growth factor 2 (FGF2; R&D Biosystems, USA). In FAot and FAits media, chicken serum was replaced by 10 µg/mL ovotransferrin (Sigma-Aldrich) or 1× Insulin-Transferrin-Selenium supplement (Gibco®), respectively. All the other components remained the same as in FAcs medium.

To prepare 3D culture media containing 0.016 % FP003, 49.2 mL of each type of medium was mixed with 0.8 mL of FP003 solution as described in the standard user manual (Nissan Chemical Industries, Ltd., Japan). These 3D media were incubated overnight at 4 °C before use. Thereafter, 3D medium containing 0.012 % or 0.010 % FP003 was obtained by mixing medium containing 0.016 % FP003 with that lacking FP003 at a ratio of 3:1 or 5:3, respectively. For cell harvesting, media were supplemented with the phosphate-buffered saline (PBS) containing 0.2 w/v % citrate (citrate/PBS).

### Measurements of physical properties

To measure sedimentation, 3D media containing 0.010 %, 0.012 %, and 0.016 % FP003 were stored in bottles and mixed with polystyrene beads with a diameter of 200–300 µm. Apparent viscoelasticity was measured using an MCR 301 rheometer (Anton Paar, Germany), a 50 mm cone plate, and a gap of 0.102 mm at 25 °C with a shear rate of 8.86 s^−1^.

### Establishment and *in vitro* culture of PGCs

cPGC lines were established by seeding 5 µL of blood obtained from the dorsal aorta of each chicken embryo at HH15–16 (Day 3 incubation, E3) into 300 µL of FAcs medium [24]. One-third of the medium was replaced by fresh medium every 2 days. Cells were sub-cultured into a larger dish in fresh medium when they became confluent. cPGCs are suspension cells, and therefore did not require trypsinization during passage.

To establish cell lines expressing fluorescent reporters, cPGCs were infected with a recombinant lentivirus harboring PGK:EGFP or VASA:tdTOMATO (S1 Fig) at a multiplicity of infection of 1. cPGCs expressing PGK:EGFP or VASA:tdTOMATO were selected by culture in the presence of 0.1 µg/mL puromycin (Gibco^®^) or 250 µg/mL G418 (Gibco^®^) for 2 weeks, respectively. Monoclonal cell lines were then established from single cells via flow cytometry (FACSAria III, BD Biosciences, USA). Viral particles were produced by co-transfecting HEK293T cells with pCMVΔR8.91, pMD.G, and a functional plasmid (pAS7w.EGFP.puro or pLAS2W-dHS4-prmVASATdTomato-pA.Pneo).

pLAS2W-dHS4-prmVASA-TdTomato-pA.Pneo was constructed by inserting the dHS4-prmVASA-TdTomato-pA cassette excised from the pPB-dHS4-prmVASATdTomato-pA plasmid, which was generated by Dr. Bertrand Pain‘s team using a 2000 bp fragment of the mouse Ddx4 promoter cloned with 5′-GGC TCT AGA GGA TCG GCC TGG GCG ACT ACA GTG-3′ forward primer and 5′-CCT TGC TCA CCA TGG GAT AGC TTC AGG TTC CTA AAA AAA AAA A-3′ reverse primer from mouse genomic DNA, into pLAS2W.Pneo from the National RNAi Core Facility (Academia Sinica, Taiwan) (S1 Fig). The procedures used to prepare viral particles were provided by the National RNAi Core Facility. Viral particle transduction and related manipulations were conducted in a BSL2 level laboratory in accordance with standard safety guidelines. All cPGC lines were maintained at 37 °C in 5 % CO_2_.

### Cell proliferation assay

Cell proliferation was assessed using a Cell Counting Kit (CCK-8/WST) (Dojindo, Japan). To determine the relative total cell number, standard curves were drawn for cPGCs at a variety of densities (1–9 × 10^5^ cells/mL) and cultured in 2D and 3D media. The cell density was plotted against absorbance at 450 nm (S2 Fig). This absorbance was measured using a Spectramax190 spectrophotometer (Molecular Devices, USA) after incubation for 4 hr at 37°C. All measurements were performed at the same time each day by mixing CCK-8 reagent and suspension media at a ratio of 1:10. The fold increase in the total cell number was calculated using the following formula: relative total cell number at Day N ÷ relative total cell number at Day 1.

### Immunofluorescence and flow cytometric analysis

Cells were washed twice and suspended in ice-cold PBS lacking Ca^2+^/Mg^2+^ (Gibco^®^). The cell suspension (5 × 10^4^ cells) was placed onto a Superfrost^™^ Plus slide (Thermo Scientific^™^, USA). After incubation for 20 min at an ambient temperature, cells had adhered to the slide and were examined by microscopy. The remaining cell suspension was analyzed by flow cytometry. For immunofluorescence staining of stage-specific embryonic antigen-1 (SSEA-1), cells attached to a slide or in suspension (5 × 10^5^ cells) were incubated overnight at 4 °C with 0.125 μg of an anti-SSEA-1 Alexa Fluor^®^ 488-conjugated antibody or a mouse IgM isotype control FITC-conjugated antibody (eBioscience, USA) in 500 μL of blocking buffer (Dulbecco‘s PBS containing 1% bovine serum albumin (Sigma-Aldrich)). Slides were washed with PBS and mounted using ProLong^™^ Gold Antifade Mountant containing DAPI (Life Technologies, USA). Images were acquired using a Leica DM2500 Optical Microscope (Leica Microsystems, Germany) equipped with a Canon EOS 7D camera (Canon, Japan). Flow cytometric analysis was conducted on a Cytomics FC500 cytometer (Beckman Coulter, USA). Data were analyzed using CXP software (Beckman Coulter).

### RNA isolation and reverse transcription PCR (RT-PCR)

Total RNA was isolated from cells using TRIzol^®^ Reagent (Invitrogen, USA) according to the manufacturer‘s instructions. Samples were resuspended in DNase/RNase-free distilled water and quantified using a NanoDrop 1000 spectrophotometer (Thermo Scientific^™^). Thereafter, 1 µg of total RNA was treated with DNase I (Invitrogen) and reverse-transcribed into cDNA using a High-Capacity >RNA-to-cDNA^™^ Kit (Applied Biosystems, USA). Expression of various genes was measured by PCR using the primer sets shown in S1 Table. In addition, expression of the housekeeping gene *GAPDH* was assessed as an internal control. The PCR mixture contained 1 U of Ultra-Pure Taq PCR Master Mix (Geneaid Biotech, Taiwan), 10 µM of each primer, 50 ng of cDNA, and ultra-pure water up to a total volume of 20 µL. The cycling conditions were 94 °C for 5 min, followed by 35 cycles of 94 °C for 20 s, 59 °C for 30 s, and 72 °C for 20 s. The PCR products were run on a 2 % agarose gel and stained with ethidium bromide.

### Gonadal migration assay

1 × 10^6^ cells of vtPGCs were centrifuged and resuspended in 100 µL of the FAot medium with 1 µL of 2.5 % Patent Blue V solution (Sigma-Aldrich, Germany). After opening a small window in each recipient egg using a mini electric driller. 1 µL of the cell suspension (around 10^4^ cells) was transferred into the dorsal aorta of each recipient embryo at HH stage 15-16 by microinjection with a sharp glass capillary (inner diameter: 30 µm), The window was sealed with Tegaderm^™^ Film (3M Health Care, USA). To observe the colonization of donor cells in embryonic gonads, the injected embryos were isolated and dissected to reveal the entire gonad one week after injection (E10). The gonads from chicken recipients were also collected and extracted genomic ADNA for further molecular analysis. The PCR analysis for the detection of tdTomato fragment in the DNA from gonads was performed by the materials and methods mentioned previously. Photographic images were obtained using an optical microscope (Leica Z16 APO, Leica Microsystems, Germany) equipped with a Canon EOS 7D camera (Canon, Japan).

### Statistical analysis

All statistical analyses were performed using GraphPad Prism 6 (GraphPad ASoftware, USA). Data are presented as mean ± standard error of the mean (SEM). p < 0.05 (calculated using a one-way ANOVA with Tukey‘s post hoc test) was considered statistically significant. For ANOVA tests with triplicate technical repeat in two independent lines at least, the different levels of significance are denoted by different symbols in the figures.

## Results

### Supplementation of FAcs medium with FP003 inhibits sedimentation

We supplemented the previously described FAcs medium with various concentrations of the functional polymer FP003 in an attempt to prevent sedimentation. Media containing and lacking FP003 are referred to as 3D and 2D media, respectively. Polystyrene beads were used to mimic cells (Fig 1A). After shaking, beads sedimented in 2D medium, but remained suspended in 3D media containing 0.010 %, 0.012 %, and 0.016 % FP003. Moreover, viscoelasticity was 0.9 mPa/s in 2D medium and 5.6 mPa/s in 3D medium containing 0.016 % FP003 (Fig 1B). These results suggest that addition of low concentrations (0.010–0.016 %) of FP003 to culture medium inhibits sedimentation but does not markedly affect viscoelasticity. cPGCs were distributed throughout 3D medium, while most settled on the bottom of the dish in 2D medium (Fig 1C).

**Fig 1.**
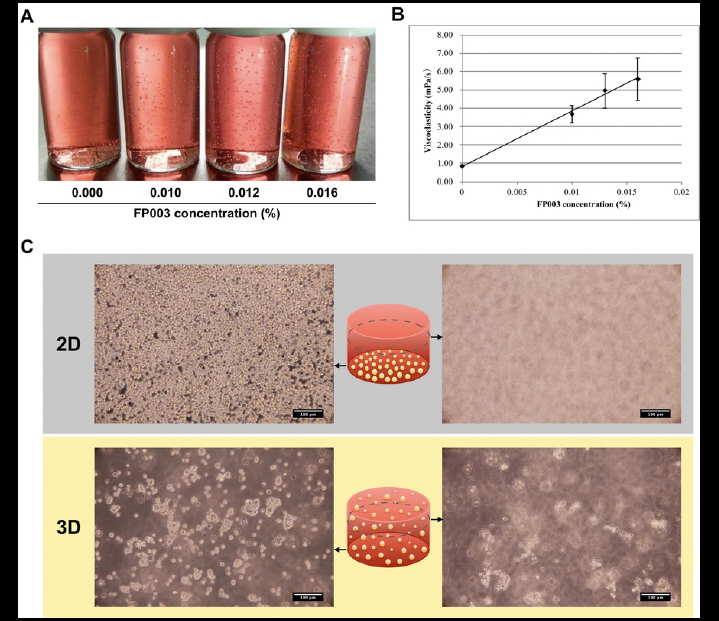
Characterization of culture medium containing FP003. (A) Sedimentation of polystyrene beads was assessed in culture medium containing various concentrations of FP003. (B) Viscoelasticity was measured in culture medium containing different concentrations of FP003. (C) Culture of cPGCs in 2D and 3D media. The images show that all cPGCs settled on the bottom of the dish in 2D medium but were distributed over all surfaces in 3D medium. Scale bar: 100 μm.

### Establishment of the optimal parameters for 3D culture of cPGCs

To establish the optimal conditions for 3D culture of cPGCs, we investigated the maximum duration of cell growth in 2D medium and 3D medium containing 0.010 %, 0.012 %, and 0.016 % FP003. cPGCs were seeded into each type of media at a density of 5 × 10^4^ cells/mL. Cell growth began to decrease after 72 hr in 2D medium, while it continued to increase up to 96 hr in 3D media supplemented with the three concentrations of FP003 (Fig 2A). After 96 hr, growth tended to decrease in all groups as cells became confluent (Fig 2A). The fold increase in the total cell number after 48 hr was significantly lower for cPGCs cultured in 2D medium than for cPGCs cultured in 3D medium containing each of the three concentrations of FP003 (Fig 2B). After 96 hr, the fold increase in the total cell number for cPGCs cultured in 3D medium containing 0.016 % FP003 was twice that for cPGCs cultured in 2D medium (Fig 2B). The fold increase in the total cell number was highest for cPGCs cultured in 3D medium containing 0.012 % FP003 after 48 hr, but for cPGCs cultured in 3D medium containing 0.016 % FP003 after 72 hr. After 96 hr, the total cell number had increased by ~6-fold for cPGCs cultured in 3D medium containing 0.016 % FP003, and the fold increase in the total cell number was significantly (p < 0.0001) higher for these cells than for those cultured in 3D media containing 0.010 % and 0.012 % FP003 (S3 Fig).

**Fig 2.**
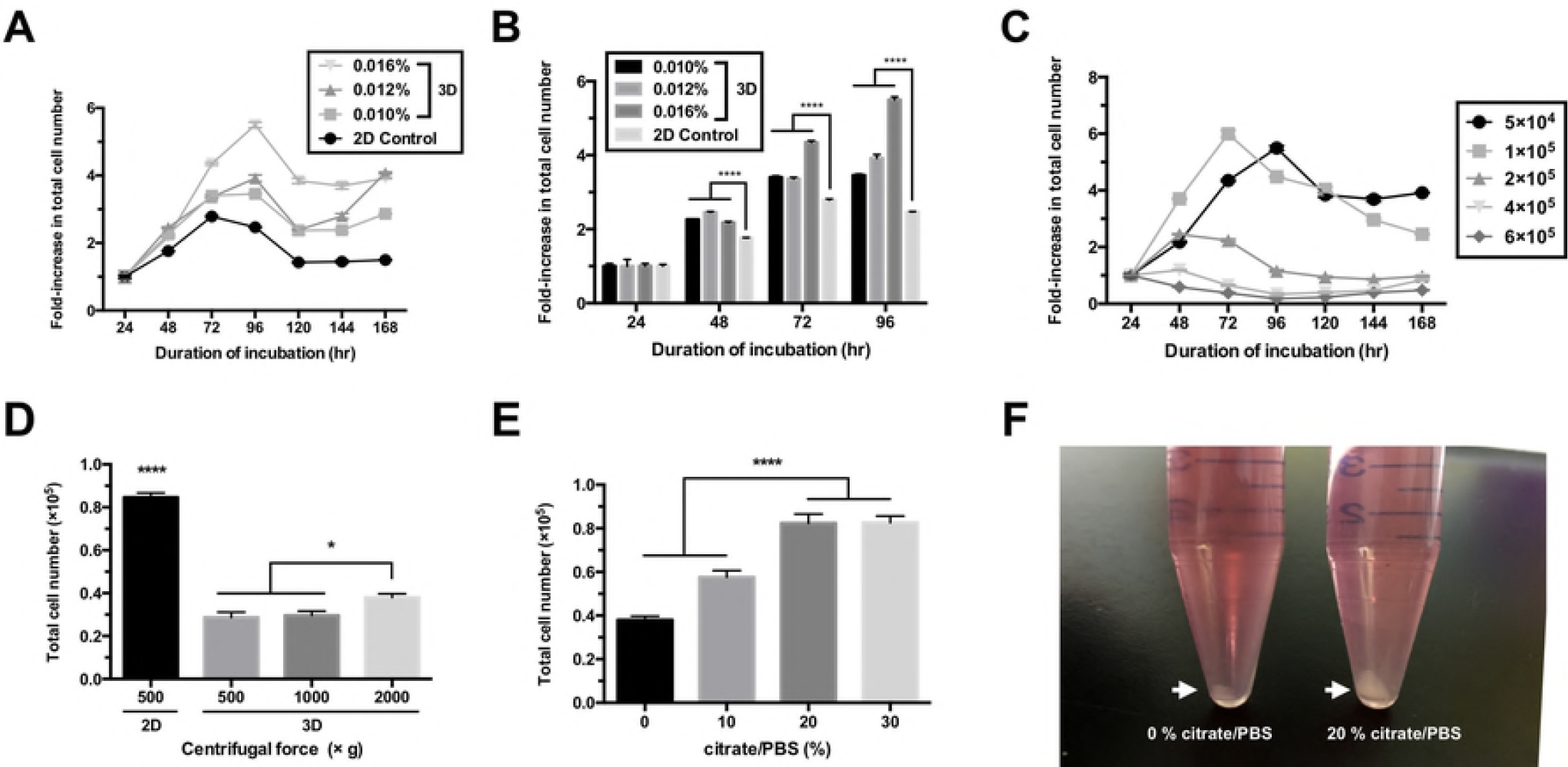
Establishment of the optimal parameters for 3D culture of cPGCs in FAcs medium. (A) Growth curves of cPGCs in 2D and 3D media over 168 hr without adding fresh medium. The 3D medium was supplemented with various concentrations of FP003. cPGCs were seeded at a density of 5 × 104 cells/mL. (B) Fold increase in the total number of cPGCs grown in 2D or 3D medium for 96 hr. (C) Growth curves of cPGCs seeded at a variety of densities and cultured in medium containing 0.016 % FP003 for 168 hr without adding fresh medium. The numbers of cPGCs per mL are indicated. (D) cPGCs cultured in 2D and 3D media were harvested by centrifugation at various forces. (E) cPGCs cultured in 3D medium were harvested by supplementing the culture with different amounts of citrate/PBS and then centrifuging the sample at 2000 × g. (F) Cells were harvested from 3D media. The cell pellets with different sizes in the two media were collected as indicated (arrows). All data are mean ± SEM. * p < 0.05; **** p < 0.0001.

To determine the optimal cell seeding density, we seeded cPGCs at five densities in FAcs medium containing 0.016 % FP003. cPGCs seeded at densities of 5 × 10^4^, 1 × 10^5^, and 2 × 10^5^ cells/mL expanded (Fig 2C). A seeding density of 1 × 10^5^ cells/mL was optimal for the proliferation of cPGCs. Using this seeding density, the total cell number was 6-fold higher at 72 hr than at 24 hr (Fig 2C). cPGCs cultured in polymer-containing 3D medium were difficult to isolate from suspension by only centrifugation when compared to those in 2D medium (Fig 2D). To harvest cPGCs in 3D medium, samples were centrifuged at 2000 × g following addition of up to 20 vol % citrate/PBS in order to dissociate polymer-ion structures. Cells were as efficiently harvested by this method as by centrifugation for 5 min at 500 × g in 2D medium (Fig 2E and 2F).

### Comparison of the growth of cPGCs between serum-containing and chemically defined media

We plated a low number (1 × 10^4^) of cPGCs in serum-containing (FAcs) or chemically defined (FAot or FAits) medium and cultured the cells for 1 week. The proliferation of cPGCs cultured in these three types of media differed at various time points (Fig 3A). The fold increase in the total cell number after 96 and 168 hr was significantly (p < 0.0001) lower in FAot and FAits media than in FAcs medium (Fig 3B). cPGCs kept proliferating over 1 week of culture in FAot medium, but not in FAits medium. The growth curve of cPGCs was similar in FAot and FAcs media.

**Fig 3.**
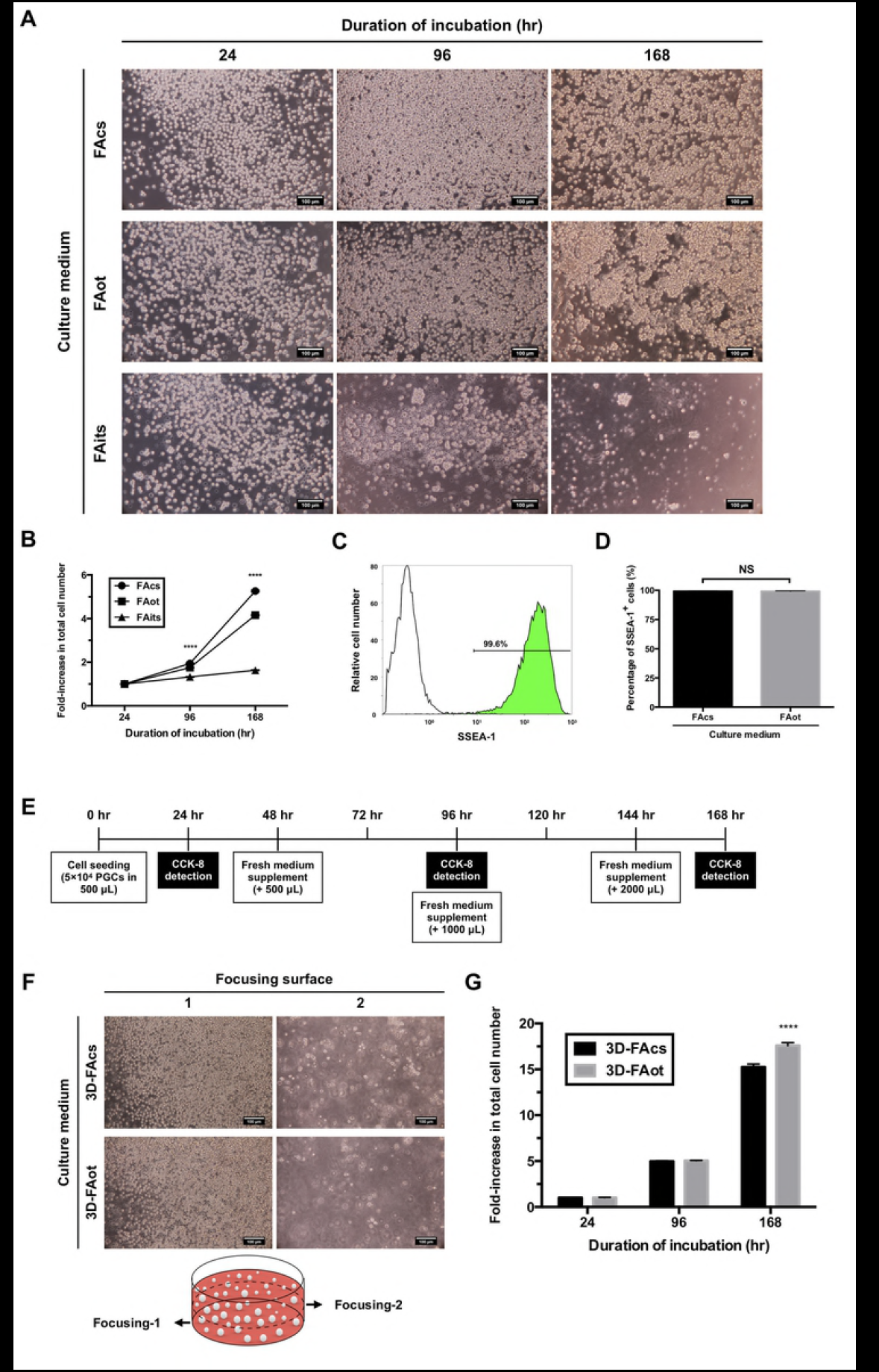
Culture of cPGCs in serum-containing or chemically defined media. (A) Images of cPGCs cultured in a serum-containing (FAcs) and chemically defined (FAot or FAits) media for 24, 96, and 168 hr. cPGCs were seeded at a density of 1 × 10^4^ cells/mL. Scale bar: 100 μm. (B) Fold increase in the total number of cPGCs after culture in each type of media for 24, 96, and 168 hr. Cell proliferation was assessed using the CCK-8 assay. Data are mean ± SEM, the statistical significance of difference among three groups was indicated. **** p < 0.0001. (C) Flow cytometric data. The number indicates the percentage of cells stained with an anti-SSEA-1 antibody (green). Isotype staining was performed as a control (white). (D) Percentages of SSEA-1+ cPGCs in FAcs and FAot media. Data are mean ± SEM. NS, not significant. (E) Timeline of the experimental protocol. (F) Images of cPGCs cultured in 3D-FAcs and 3D-FAot media. The images were acquired by focusing on one of two surfaces, which are indicated by arrows in the cartoon. Scale bar: 100 μm. (G) Fold increase in the total number of cPGCs upon culture in 3D-FAcs and 3D-FAot media for 24, 96, and 168 hr. Data are mean ± SEM. **** p < 0.0001.

cPGCs were cultured in FAcs or FAot medium for 1 week, subjected to immunofluorescence staining of the pluripotent cell surface marker SSEA-1, and analyzed by flow cytometry. SSEA-1 staining was significantly more intense than isotype antibody staining. More than 99 % of cPGCs cultured in FAcs or FAot medium were SSEA-1+, and there was no significant difference between the two groups (Fig 3C and 3D).

### Expansion of cPGCs in 3D medium containing or lacking serum

We further investigated the proliferation of cPGCs in FAcs and FAot media containing FP003. To this end, 5 × 10^4^ cPGCs were suspended in 0.5 mL of FAcs or FAot medium containing 0.016 % FP003 (3D-FAcs and 3D-FAot media, respectively), and the same volume of fresh medium was added every 2 days (Fig 3E). The CCK-8 assay was used to determine the total cell number after 24, 96, and 168 hr. cPGCs remained distributed throughout both 3D-FAcs and 3D-FAot media (Fig 3F). Moreover, the fold increase in the total cell number after 168 hr was significantly larger in 3DFAot medium (17.6-fold) than in 3D-FAcs medium (15.2-fold) (Fig 3G).

### Maintenance of PGC characteristics upon long-term 3D culture

To determine whether stem cell properties were maintained upon long-term culture in 3D media, we established cPGCs that expressed the germ cell-specific reporter VASA:tdTOMATO (vtPGCs) to monitor germline identity in real-time. vtPGCs stably expressed the germline reporter over 4 weeks of culture in 3D-FAcs and 3D-FAot media (Fig 4A). The pluripotency of vtPGCs cultured in 3D media was investigated by performing immunofluorescence staining of SSEA-1. SSEA-1 was detected on the surface of vtPGCs (Fig 4B). Flow cytometry demonstrated that the percentages of vtPGCs positive for tdTOMATO and SSEA-1 were higher than 97 % following culture in 3D-FAcs or 3D-FAot medium for 4 weeks and did not markedly differ between the two groups (Fig 4C). Reverse transcription PCR (RT-PCR) analysis demonstrated that germline-specific (*DDX4* and *DAZL*) and pluripotency-associated (*POUV*/*OCT4* and *NANOG*) genes, as well as *PRDM1* and *PRDM14*, which encode critical regulators in PGCs, were highly expressed in pre-cultured and cultured cPGCs and vtPGCs, but were not expressed in chicken embryonic fibroblasts, which were used as a somatic cell control (Fig 4D). Moreover, vtPGCs stably expressed tdTOMATO upon long-term culture in 3D media. In addition, the gonadal migration as the key function of cPGC was exhibited after the transplantation of long-term cultured vtPGCs (S4 Fig). The vtPGCs long-term cultured in 3D condition could successfully colonized in the gonads of recipients following the migration through circulation (S4 Fig).

**Fig 4.**
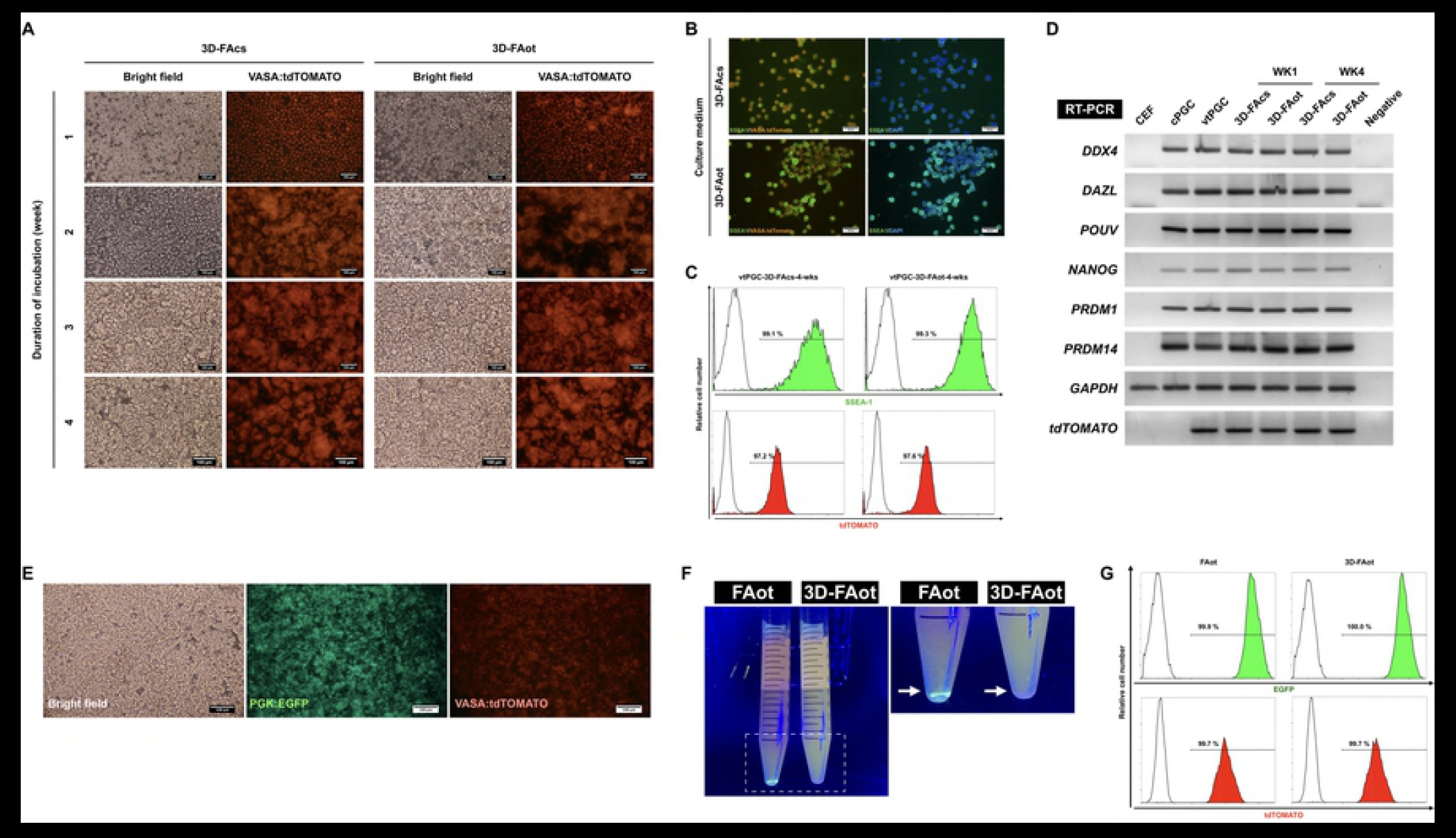
Characterization and ectopic protein expression of cPGC lines cultured for a long term in 3D media. (A) Images showing the proliferation of vtPGCs and their expression of the germline-specific reporter tdTOMATO over 4 weeks of culture in 3D-FAcs and 3D-FAot media. Red labeling corresponds to tdTOMATO. Scale bar: 100 μm. (B) Immunofluorescence staining of SSEA-1 in vtPGCs cultured for 4 weeks in 3D-FAcs and 3D-FAot media. Green, red, and blue staining corresponds to SSEA-1, tdTOMATO, and DAPI, respectively. Scale bar: 50 μm. (C) Flow cytometric analysis of SSEA-1 and tdTOMATO expression in vtPGCs cultured for 4 weeks in 3D-FAcs and 3D-FAot media. The percentage of positively labeled vtPGCs is indicated in each graph. cPGCs were stained with mouse IgM isotype antibodies as a control. (D) RTPCR analysis of the expression of pluripotency-related and germline-specific genes in vtPGCs cultured in 3D-FAcs and 3D-FAot media. *GAPDH* was used as an internal control. CEF, chicken embryonic fibroblast. (E) DuotonePGCs expressed EGFP and tdTOMATO. Scale bar: 100 μm. (F) Sedimentation of DuotonePGCs was assessed in FAot and 3D-FAot media. DuotonePGCs were largely precipitated in FAot medium and evenly distributed in 3D-FAot as the arrows indicated under the fluorescent photography. (G) Flow cytometric analysis of EGFP and tdTOMATO expression in duotonePGCs cultured in FAot and 3D-FAot media. The percentage of positively labeled cells is shown in each graph.

### Ectopic expression of recombinant fluorescent proteins in cPGC lines upon culture in 3D-FAot medium

We attempted to produce recombinant fluorescent proteins in cPGCs harboring PGK:EGFP and VASA:tdTOMATO (duotonePGCs) via culture in 3D-FAot medium (Fig 4E). By fluorescent photography, duotonePGCs apparently showed an even distribution in 3D-FAot, compared to that cells were sedimented in FAot medium after static settlement for 20 minutes (Fig 4F). Moreover, Flow cytometric analysis demonstrated that almost all cells expressed EGFP and tdTOMATO upon culture for 1 week in FAot and 3D-FAot media, and the percentages of positive cells did not markedly differ between the two groups (Fig 4G). These results indicate that culture in 3D-FAot medium supports the growth of the cPGC lines and the production of recombinant fluorescent proteins.

## Discussion

Various strategies have been developed for 3D cell culture, including those that use scaffolds, the hanging drop technique, and polymers. The polysaccharide low-acyl gellan gum has been widely used to form double helical structures with cations [25]. These structures inhibit sedimentation of cultured cells and cellular spheroids and can be used to develop a 3D culture system for various purposes, including drug screening [26], accelerated differentiation [27], and production of stem cells for clinical applications [1, 2].

FP003, the functional polymer used in the present study, contains a small amount of gellan gum and thus forms structures that prevent cell sedimentation. cPGCs must be cultured in medium containing a low concentration of calcium (< 0.15 mM) to prevent their aggregation [24]. This may influence the interaction between polymers and calcium ions and thus the formation of structures that inhibit sedimentation. Fortunately, such structures still formed in culture medium containing a reduced level of calcium ions and other cations. One reason inorganic salts are added to cell culture media is to adjust the osmotic pressure, and many types of media contain a moderate level of cations. Cells did not sediment in the FP003-containing media used in this study, despite the relatively low concentration of calcium, suggesting that FP003 are useful for 3D cell culture in this situation. On the other hand, the efficiency of cell harvesting from media containing gellan gum (FP001) is low. To recover single cells or spheroids from FP001-containing medium, the cell suspension must be diluted several fold with fresh medium or PBS. FP003 contains less gellan gum than FP001. Therefore, we evaluated the efficiency of cell harvesting from medium containing FP003 using citrate as a chelating agent. Addition of more than 20 vol % citrate/PBS to the cell suspension improved the efficiency of cell harvesting from FP003-containing medium. Citrate chelated with several types of cations in the culture medium, and consequently the 3D polymer network was dissociated.

Glycosylation of avian-derived proteins for therapeutic purposes was recently discussed. Eggs are considered an ideal platform for recombinant protein production, and the ovalbumin promoter shows a robust and specific expression ability in oviducts. Therefore, numerous studies have attempted to produce recombinant proteins using transgenic hens [13, 16, 17]. In addition, oviduct bioreactor also presents the easiness in transgenic animal production and husbandry, as well as the recombinant protein purification. Therefore, transgenic recombinant proteins can be more easily produced in oviduct bioreactors than in mammary gland bioreactors [28]. Despite that, animal bioreactors are more sophisticated in operation than cell-mediated production systems. Moreover, the animal safety is always an issue criticized for pharmaceutical purpose. However, compared to mammalian cells, only a small number of avian cell types can be cultured *in vitro*. Similar to chicken embryonic stem cells [29], cPGCs [24, 30] are pluripotent and can divide indefinitely when cultured under suitable conditions. We optimized the conditions for 3D culture of cPGCs in chemically defined media. These media contained Activin A, FGF2, and insulin or ovotransferrin. cPGCs could be cultured for a long period of time in a chemically defined medium without loss of cellular properties, and their proliferation was higher in 3D media than in 2D media. This stable 3D culture system displays a scalable production in cPGCs without requiring stirred tank bioreactor and associated equipment. In addition, genetic modifications are easily introduced into cPGCs. With the development in novel strategies, the efficient transgene insertion and even the precision modification in genome have been proven to obtain in this cell type [31–34]. To establish fluorescent protein-expressing cPGC lines, we selected single cells for amplifying to each clonal cell lines via fluorescence-activated cell sorting after viral transduction. These cells ectopically expressed fluorescent proteins. Our results indicate that genetically modified cPGCs can be expanded on a large scale using this 3D culture system and used to produce various recombinant proteins with therapeutic uses. Thus, cPGCs are not only useful for the production of genetically modified chickens due to their germline competence but are also a potential platform for recombinant protein production.

In summary, we showed that 3D-FAot medium can be used for long-term culture of cPGCs. cPGCs remained distributed throughout 3D-FAot media without stirring (Fig 5). Our system makes efficient use of culture space and resources. The total number of cPGCs increased by ~17-fold upon culture in 3D-FAot medium for 1 week. Moreover, the characteristics and functions of cPGCs were maintained upon culture in 3D-FAot for 1 month, and these cells stably expressed recombinant fluorescent proteins from the expression cassettes. Taken together, this 3D cell culture technique is applicable for the large-scale production of cPGCs and other applications.

**Fig 5.**
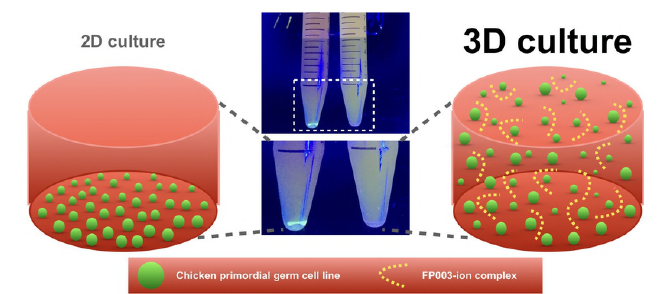
Graphical summary of the 3D chemically defined culture system for cPGC line by using FP003, and the comparison with the 2D condition.

## Acknowledgements

This work was supported by the Council of Agriculture, Executive Yuan, Taiwan (grant numbers 106AS-2.2.2-AD-U1(Z) and 107AS-2.2.2-AD-U1 awarded to SCW) and by ANR, France (grant number CRB-ANIM-ANR-11-INBS-0003 awarded to BP). We thank Mr. Nobutomo Tsuruzoe, Dr. Masato Horikawa, Ms. Chun-Yun Gu, and Ms. Charlene Hung (Nissan Chemical Industries, Ltd.) for assistance and the National RNAi Core Facility, Academia Sinica, Taiwan for providing recombinant lentiviruses and services. Dr. Hsinyu Lee from National Taiwan University kindly allowed us access to a BSL2 level laboratory.

## Supporting information

**S1 Fig. Fluorescent protein-expressing vectors used to establish vtPGCs and duotonePGCs.** (A) Diagram of the cassette containing PGK:EGFP and associated plasmid features in the lentiviral vector. The total length of the fragment is 3501 bp. (B) Diagram of the cassette containing VASA:tdTOMATO and related plasmid features. The total length of the fragment is 9195 bp. (C) Images of cPGCs expressing these fluorescent reporters. Scale bar: 50 μm.

**S2 Fig. Standard curves were generated by plotting relative absorbance at 450 nm.** As determined by the CCK-8 assay, against the seeding density of cPGCs cultured in 2D or 3D medium. The formula and R-square value are provided next to each curve. Data are the mean. Each curve was generated using three replications.

**S1 Table.**
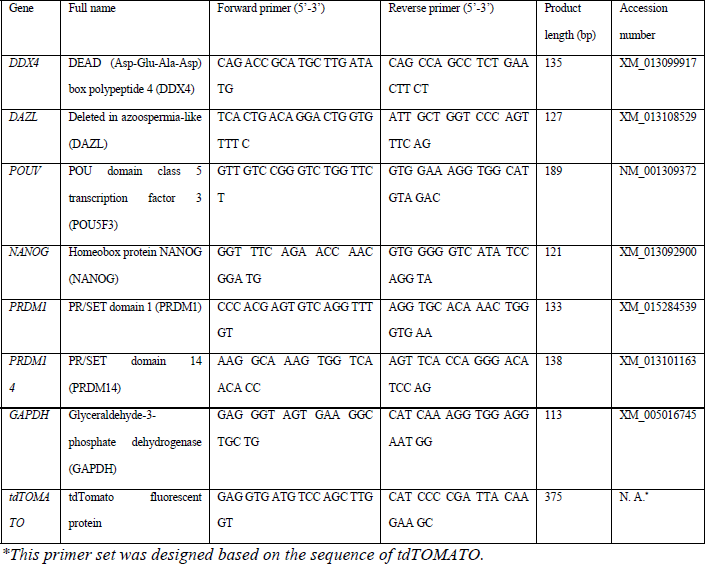
Primer sets used for RT-PCR.

**S3 Fig. Fold increase in the total number of cPGCs grown in 3D medium containing various concentrations of FP003 for 96 hr.** All data are mean ± SEM. * p < 0.05; *** p < 0.001; **** p < 0.0001

**S4 Fig. Gonadal homing migration of vtPGCs after 3D culture for 4 weeks.** (A) The detection of tdTomato gene fragment in chicken embryonic gonads with or without the transplantation of 3D cultured vtPGCs by the PCR for a specific template. The template sized 375-bp represented the positive PCR product of tdTomato gene. (B) After PGC transplantation at E3, photographs indicated the E10 embryonic gonad with the colonization of the exogenic vtPGCs undergone the 4-week-culture in 3D-FAcs or (C) 3D-FAot medium. Scale bar: 1 mm (upper); 0.1 mm (below).

